# Mechanism of the Sequential Trajectory of Allosteric Ring Closure in Human Chaperonin TRiC

**DOI:** 10.64898/2026.10.20.760597

**Authors:** Yanyan Zhao, Michael F. Schmid, Wah Chiu

**Author notes:** Correspondence (Y.Z.).

## Abstract

Eukaryotic chaperonin TRiC assists in the folding of approximately 10% of the cytosolic proteome through an ATP-driven chamber closure cycle, yet the underlying closure mechanism has remained elusive. Here, using cryo-EM and deep learning analyses, we resolve, at near-atomic resolution, the temporal engagement sequence of the eight distinct subunits during human TRiC ring closure triggered by ATP binding. TRiC closes its chamber through the stepwise extension of an apical helical network among subunits around the ring, coordinated with sequential rotations of the equatorial domains. Each subunit responds differently to ATP, rendering the overall motion asymmetric and ordered. This work delineates the complete structural trajectory of allosteric propagation during ATP-binding-induced, rather than hydrolysis-induced, ring closure, and establishes a mechanistic framework linking ATP-driven conformational dynamics to substrate handling.

## Introduction

The human chaperonin TRiC plays a central role in maintaining cytosolic proteostasis by assisting the folding of a broad array of essential client proteins, including actin, tubulin, and many regulators involved in the cell cycle and signaling pathways^1,2^. Its ability to manage the folding of such a diverse range of proteins arises from its highly sophisticated and uniquely complex structure. Unlike bacterial chaperonins, which typically consist of identical subunits arranged in symmetrical rings, eukaryotic TRiC (∼1MDa) is composed of eight distinct paralogous subunits (CCT1–CCT8) assembled into two stacked rings, each containing one copy of every subunit in a strictly defined order^3–5^ (Figure S1). This hetero-oligomeric arrangement creates an asymmetric folding chamber, endowing TRiC with specialized substrate recognition capabilities^6–10^.

TRiC operates through an ATP-dependent conformational cycle. In its open state, TRiC binds unfolded or partially folded substrates. Upon ATP binding and hydrolysis, the chaperonin undergoes a large conformational change to form an enclosed chamber that isolates the substrate from the cytoplasm. TRiC assists substrate folding both during its ATP-driven ring closure pathway and within its fully closed folding chamber. For example, previous study has shown that the open TRiC complex establishes a specific actin topology that promotes productive folding. ATP binding then induces asymmetric ring closure, progressively releasing actin segments and priming the substrate for final folding once fully encapsulated within the chaperonin cavity^11^. TRiC ring closure dynamics are mechanistically coupled to the ordered sequence of substrate folding.

Each TRiC subunit shares a conserved domain architecture: an equatorial ATP-binding domain, an apical lid domain, and an intermediate domain that connects the two and mediates conformational changes. From open to closed conformation, the equatorial domains of subunits rotate outward as apical domains rotate inward to form a lid over the chamber. Yet the ring closure mechanism remains unknown, leaving unresolved how eukaryotic TRiC—a hetero-oligomeric 16-subunit complex—closes its two rings, whether in a sequential, subunit-by-subunit manner or through simultaneous movement of multiple subunits, in response to ATP. Understanding this process could reveal how inter-subunit communication coordinates conformational transitions, ensuring efficient and specific protein folding.

To address the question of ring closure mechanism in eukaryotic chaperonin TRiC, here we use cryogenic electron microscopy (cryo-EM) to capture physiologically relevant subunit responses in human TRiC in the presence of ATP. We then apply a deep learning-based heterogeneity analysis to probe conformational variability of ATP-bound TRiC. This approach reveals eight high-resolution intermediate structures along the sequential allosteric pathway of chamber closure.

These ATP bound structures demonstrate that ATP binding alone, rather than subsequent hydrolysis, is sufficient to drive sequential ring closure, coupling two parallel conformational switch mechanisms: (i) the stepwise formation of the apical protrusion helix network and (ii) the progressive rotation of subunit equatorial domains. Our study, by resolving the detailed structural trajectory of TRiC’s ring closure, provides a mechanistic framework for future studies of investigating how TRiC ring closure translates into substrate handling.

## Result

To investigate the mechanism of TRiC ring closure in the presence of ATP, purified human TRiC complexes were mixed with 50 μM ATP and flash-frozen approximately 20 s later to capture physiologically relevant intermediates induced by ATP binding^12,13^. High-resolution cryo-EM was used to visualize these conformational states along the ATP-driven closure pathway. To resolve the complete continuum of conformations within the ensemble, we combined conventional 3D classification with iterative deep learning–based embedding to analyze 1.8 million highly heterogeneous particle images. This approach enabled homogeneous convergence within individual subsets corresponding to each conformation and revealed many transient states that conventional analysis alone failed to detect (Figures S2 and S3).

The full dataset captures TRiC particles spanning relatively open to closed states. The two rings of each complex generally move in concert but are not perfectly synchronized, with most variability localized to the apical domains of equivalent subunits across the two rings. Our deep-learning heterogeneity analysis reveals a broad conformational continuum of ATP-induced ring closure (Figures S3 and S4), but presenting every observed intermediate would obscure the underlying mechanism. Therefore, we focused on particles in which the two rings adopt closely similar conformations to (i) improve reconstruction resolution and (ii) simplify interpretation by describing subunit motions primarily in a single representative ring, thereby avoiding overly intricate structural details that might obscure the primary insights regarding allosteric propagation during closure. From these data we identify eight major near-atomic resolution states (R1–R8) as the most informative representatives of this continuum, and propose their ordering along the ring-closure pathway (Figure 1, Figures S5–S8). R1-R5 represent the five major checkpoints where the ring remains in a generally open state (31.6% of the 1,630,427 open-state particles), while R6-R8 represent later-stage checkpoints with nearly or fully closed rings (221,070 closed-state particles). The majority of remaining 68.4% of open-state particles represent continuous intermediate states between these checkpoints and are not explicitly described in the manuscript (Figures S3 and S4).

**Figure 1.**
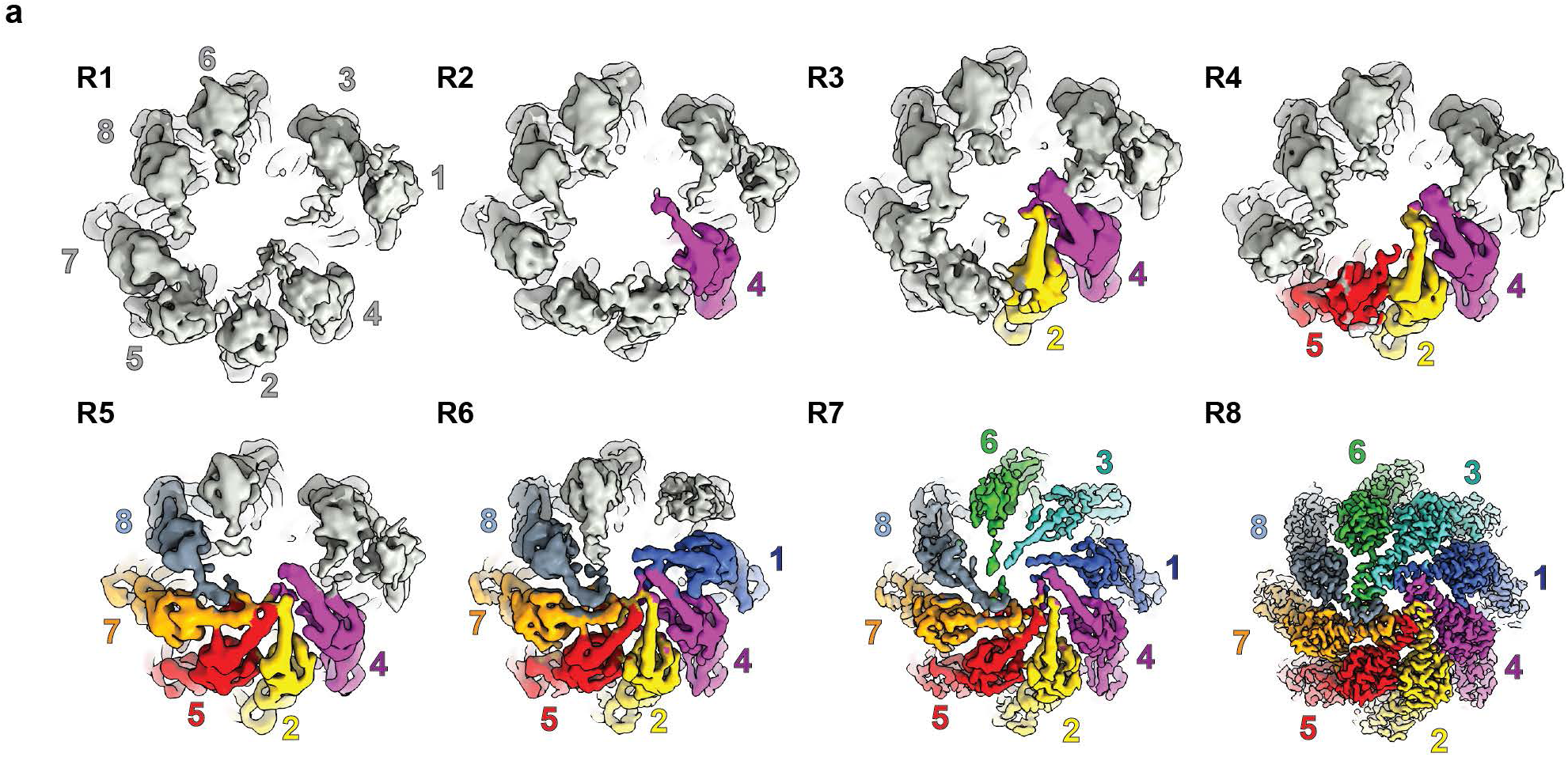
The eight cryo-EM reconstructions of human chaperonin TRiC in the presence of ATP.

### ATP-Bound Pre-Closure State of TRiC

A subset of TRiC particles contributes to a highly open conformational state of TRiC, resolved at 4 Å (R1 state) (Figure 1, Figures S6, S7). In this state, the protrusion helices in the apical domains of the CCT subunits angle outward, away from the central axis of the chamber (Figure 2d). The conformations of subunits in both rings exhibit a high degree of structural similarity (Figures S7, S8). This R1 open form closely resembles the prefoldin-receiving *apo* human TRiC conformation observed without ATP (EMDB-32823)^7^, and the yeast TRiC bound with AMPPNP (EMDB-9541)^14^. The superposition of these maps reveals no major structural differences (Figure S9). Notably, in R1, ATP is bound to all CCT subunits, with CCT6 and CCT8 additionally accommodating a water molecule adjacent to the bound ATP (Figure S10). This water molecule, positioned by the aspartic acid of helix 12 in the intermediate domain, highlights the presence of a tightly closed ATP pocket (Figure 2e, Figure S10). These findings suggest that in many TRiC particles, rapid and uniform ATP loading across TRiC precedes any significant conformational changes by individual subunits. Yeast TRiC-AMPPNP (EMDB-9541), previously interpreted as the final ATP-bound state, thus actually corresponds to the early human TRiC-ATP intermediate (R1 state) before ring closure begins. AMPPNP does not faithfully recapitulate the structural consequences of ATP binding in TRiC, contrary to the common assumption in previous studies.^14,15^

**Figure 2.**
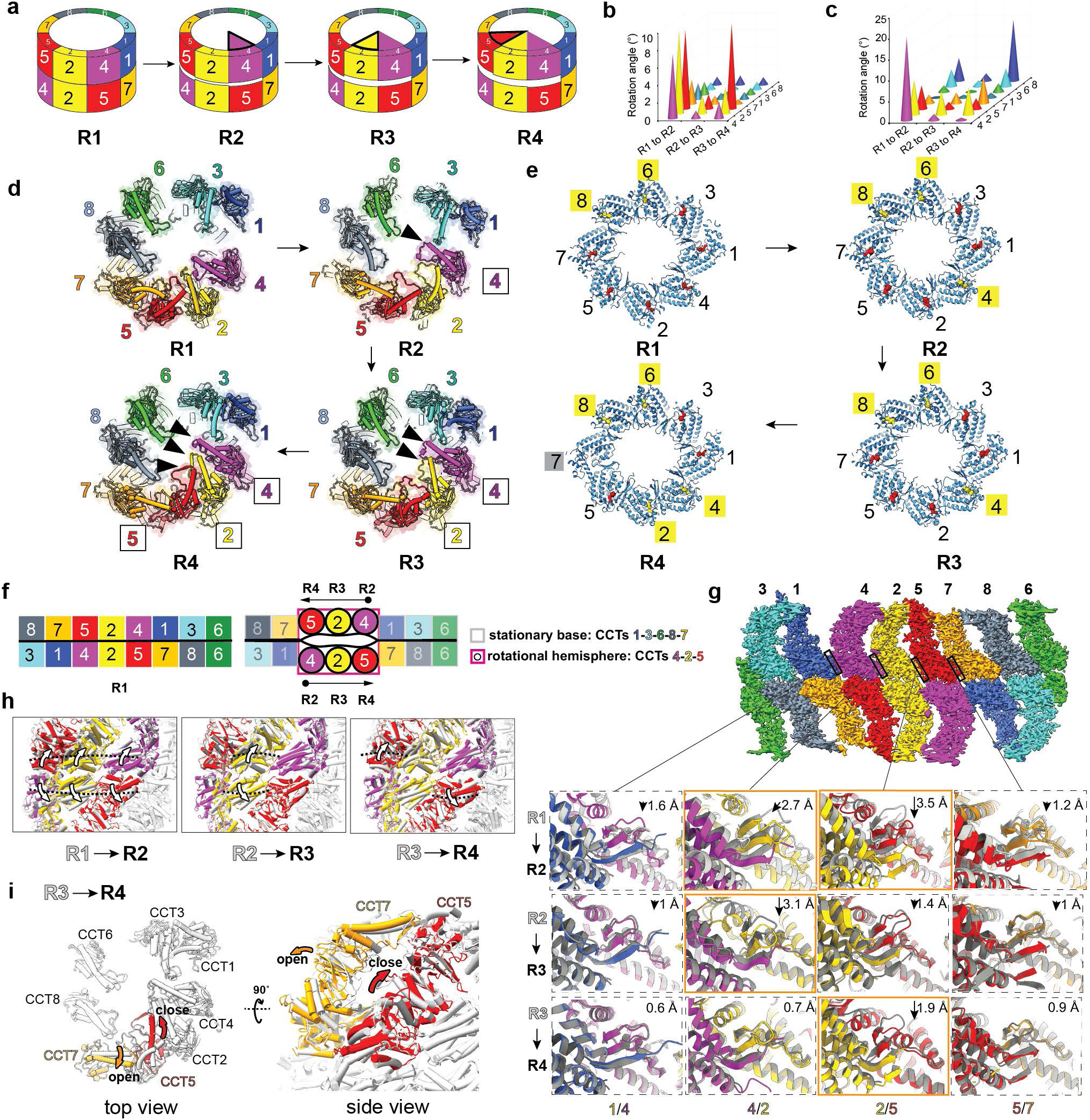
The closure of hemisphere CCTs 4-2-5. a. The schematic of sequential closure of CCTs 4-2-5 from R1 to R4, describing the initiation and extension of apical protrusion network, and the inter-ring seperation at CCTs 4-2-5. b. The histogram quantifying the equatorial domain rotations through states R1, R2, R3, R4. **c**. The histogram quantifying the intermediate domain rotations towards equatorial domain, indicates the sequential primary ATP binding response, **d.** The top view of structural models of R1-R4, showing the initiation and sequential extension of apical protrusion network by CCTs 4-2-5. **e**. The ATPMg^2+^H2O (yellow) and ATPMg^2+^ (red) distribution in R1, R2, R3, R4., and CCT7 in R4 lack ATP binding, **f**. The unwrapped arrangement of CCTs in TRiC in states R1 and R2, R3, R4, demonstrating inter-ring separation due to concerted equatorial domain rotations in CCTs 4-2-5 across the rings. The arrows indicates the primary ATP binding responder in each state of R2, R3, R4. **g.** The inter-subunit betasheet transitting the allostery across four states (R1 to R4). **h**. The superimposed models shows gradual inter-ring separation by equatorial rotation at CCT4-2-5 across R1, R2, R3, R4. **i**. The top and side view showing the apical separation between CCT5 and CCT7 in state R4.

### Closure Initiates at CCT4 and Progresses through CCT2–CCT5

In the R2 state (4.1 Å), CCT4 adopts a closed conformation ahead of the other subunits, with its apical domain and protrusion helix tilting inward toward the chamber’s central axis. Examination of all latent space volumes reveals that CCT4 is the only subunit that initially anchors the protrusion helix and therefore the first to close. This likely arises from its greater conformational freedom, afforded by the large gap between CCT4 and CCT1. In R3 (4.0 Å), CCT2 follows, positioning its protrusion helix alongside CCT4. The stronger density of both helices suggests increased structural stability or reduced flexibility. In R4 (3.9 Å), CCT5 closes next, aligning its protrusion helix with that of CCT2 (Figure 1, 2a, 2d, Figures S6-S8, Videos S1, S2, S3).

ATP binding shows differential occupancy among subunits (Figure 2e, Figure S10, Table S1). In R2, ATP is bound to all CCT subunits, with a water molecule near the putative nucleophilic site of CCT4, CCT6, and CCT8. In R3, ATP remains bound to all subunits except CCT7, which shows partial occupancy (Figure S10, Table S1). In R4, CCT7 lacks ATP entirely, while the water molecule is also observed in CCT2 (Figure S10, Table S1). This indicates that CCT7 has the weakest ATP-binding affinity among the subunits. The CCT subunits exhibit distinct conformational responses to ATP binding (Figure 2c, Figure S12). CCT4 closes its nucleotide-binding pocket fully in a single step, with its intermediate domain rotating downward by 20° toward the equatorial domain at R2. CCT2 closes gradually, rotating 8.4° at R2 and an additional 5.7° at R3. CCT5 shows only a slight 5.6° rotation of its intermediate domain at R2 and remains essentially stationary. Therefore, the extent of ATP pocket closure differs across CCT subunits within each state and progresses gradually from R1 to R4 as the lid of CCTs 4-2-5 closes.

ATP binding triggers a downward rotation of the intermediate domains, which in turn drives rotation of the equatorial domains and separation of inter-ring subunits (Figure S1). Compared to the R1 open state, the equatorial domains of CCT4, CCT2, and CCT5 undergo substantial rotations in R2 (7.4°, 9.6°, and 9.1°, respectively) and continue to rotate in R3 (0.9°, 2.1°, and 1.7°). In R4, CCT5 shows a pronounced additional rotation of 9.8°. In contrast, the equatorial domains of CCT1, CCT3, CCT6, CCT8, and CCT7 remain largely stationary throughout these states (Figures 2b, 2f, 2h, Figures S13a, S13b, S13c, S14a, Videos S1, S2, S3).

Intra-ring contacts between equatorial domains are mediated by β-sheets formed between the stem loop of one subunit and the C-terminal strand of an adjacent subunit (Figure S1). These β-sheets stabilize the ring architecture and facilitate allosteric communication. From R1 to R2, the β-sheets between CCT4–CCT2 and CCT2–CCT5 shift downward by 2.7 Å and 3.5 Å, respectively, reflecting the rotations of CCT4, CCT2, and CCT5 (Figure 2h, 2g). The β-sheets between CCT1–CCT4 and CCT5–CCT7 show only minor shifts despite substantial rotations of CCT4 and CCT5. In R3, the CCT4–CCT2 β-sheet shifts downward by 3.1 Å, while other interfaces remain largely unchanged (Figure 2g). In R4, the CCT2–CCT5 β-sheet shifts downward by 1.9 Å to accompany CCT5’s rotation, with all other β-sheet interfaces remaining stable (Figure 2g).

Shifts in β-sheet positions correlate with the equatorial domain rotations of CCT4, CCT2, and CCT5, while interfaces among the stationary subunits remain largely unchanged. Within the stationary hemisphere (CCT1, CCT3, CCT6, CCT8, CCT7), inter-subunit β-sheets act as a stable scaffold, with the CCT1–CCT4 and CCT5–CCT7 interfaces serving as pivot points that anchor the dynamic subunits (Figure 2g). Meanwhile, the β-sheets between CCT4–CCT2 and CCT2–CCT5 coordinate the concerted rotations of the moving subunits. The reverse-aligned arrangement of CCT5, CCT2, and CCT4 across the cis and trans rings (4–2–5 above 5′–2′–4′) ensures synchronized inter-ring rotation (Figure 2f).

In R3, CCT7 adopts a slightly more open conformation, with its intermediate domain rotating – 2.6° away from the equatorial domain, consistent with partial ATP occupancy in its nucleotide-binding pocket (Figures S10, S12). In R4, CCT7 opens further, with the intermediate domain rotating –6.3°, reflecting the complete absence of ATP (Figure 2e, Figures S10, S12). During this transition, ATP binding in CCT5 drives inward movement of its apical domain, while the absence of ATP in CCT7 causes its apical domain to shift outward, fully disrupting the apical interaction between CCT7 and CCT5 that is maintained from R1 to R3 (Figures 2i, 2d, Figures S7, S8, Video S3).

Throughout states R2, R3, and R4, the intermediate domain of CCT5 remains largely stationary relative to its equatorial domain, showing minimal movement (Figure 2c, Figure S12). Notably, helix 12 of CCT5’s intermediate domain occupies the same position as in the fully closed TRiC state (R8). In R4, however, disruption of the apical interaction with CCT7 is accompanied by a large rotation of CCT5’s equatorial domain, indicating that its closure had been partially restricted by the apical contact with CCT7 in earlier states.

Together, the R1–R4 states demonstrate that differential responses to ATP binding by individual CCT subunits drive asymmetric ring closure within the ring (Figures 2a, 2f). In the dynamic hemisphere, comprising CCT4, CCT2, and CCT5, closure occurs sequentially: CCT4 initiates by anchoring its apical protrusion helix toward the central vertical axis, followed by CCT2 and CCT5, which extend the protrusion network. Throughout this transition, the apical domains of CCT4, CCT2, and CCT5 tilt inward concurrently but to different extents, while their equatorial domains undergo differential outward rotation.

Under our experimental conditions (1 µM TRiC, corresponding to 16 µM total CCT subunits, mixed with 50 µM ATP—an approximately 3-fold molar excess over total subunits), and with the sample flash-frozen immediately after ATP addition to capture the intermediate, many particles achieve full ATP saturation across all 16 subunits. However, a fraction exhibits incomplete ATP occupancy, most notably at CCT7 of R4. This selective substoichiometry indicates that CCT7 has the lowest ATP-binding affinity among the eight CCT subunits. CCT7 likely restricts the early inward tilting of CCT5’s protrusion helix through its apical interaction, ensuring coordinated, asymmetric closure of the CCT ring (Videos S1, S2, S3).

### Sequential Closure at the CCT7–CCT1 Transitional Interface

In state R5, resolved at 5.5 Å, both CCT5 and CCT7 orient their protrusion helices toward the chamber’s central axis. Together with CCT4 and CCT2, they form a stable protrusion network (Figures 1, 3b, Figures S6, S7, S8, Video S4). At this stage, a β-sheet barrel beneath the protrusion network at the chamber apex begins to assemble through interactions of the apical loops from CCT7 and CCT5, as indicated by the observed density (Figure 3d). The protrusion helix of CCT8, though weaker in density, lies adjacent to that of CCT7 (Figure 1).

**Figure 3.**
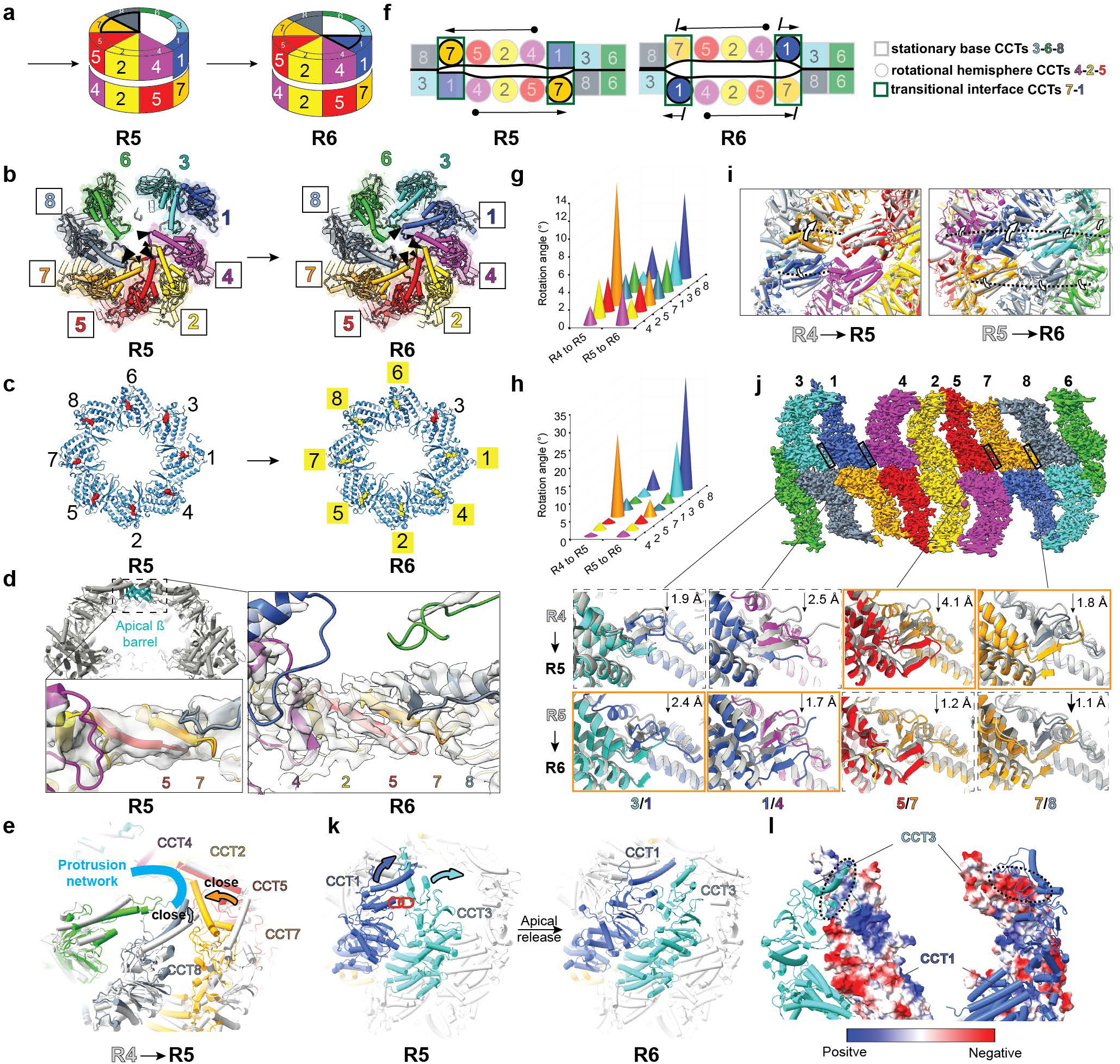
The sequential closure of CCTs 7-1. **a**. The schematic of sequential closure of CCTs 7-1 in R5 and R6. **b**. The top view of structural models of R5 and R6, showing the sequential extension of apical protrusion network by CCTs 7-8-1. **c**. The ATPMg^2+^H2O (yellow) and ATPMg^2+^ (red) distribution in R5, R6. **d.** The initiation of apical β-barrel in R5 and extensiton in R6. **e**. The CCT7 recruits CCT8 into the protrusion network in R5. **f.** The unwrapped arrangement of CCTs in TRiC in states R5 and R6, demonstrating the sequential response to ATP binding by CCT7 and CCT1. **g**. The histogram quantifying the equatorial domain rotation across R4, R5, R6. **h.** The histogram quantifying intermediate domain rotation towards equatorial domain across R4, R5, R6. **I.** The inter-ring separation by equatorial rotation at CCT7-1 across R4, R5, R6. **j**. The inter-subunit betasheet transitting the allostery across R4, R5, R6. **k**. The titled top view showing the apical separation between CCT1 and CCT3 in state R6. I. The complementary electrostatic distribution at the CCT1/3 apical interacting interface.

All CCT subunits in R5 contain ATP in their nucleotide-binding pockets; however, no water molecules are resolved near the ATP, likely due to limited resolution (Figure 3c, Figure S10). Relative to R4, the intermediate domain of CCT7 rotates downward by 23° toward its equatorial domain, closing the nucleotide-binding pocket (Figure 3h, Figure S12). The equatorial domain simultaneously rotates 13.8° within the ring, while the apical domain tilts toward CCT8 (Figure 3g, 3i, 3e, Figure S13d). As CCT7 approaches CCT8, the latter begins to engage with the protrusion helix network, even though its equatorial domain has not yet undergone significant rotation (Figure 3e, Video S4).

Closure of CCT7 destabilizes the previously stationary scaffold formed by CCT1, CCT3, CCT6, CCT8, and CCT7 in R1–R4, triggering rotational movements across the equatorial domains of all subunits except CCT3 (Figure 3g, Figure S13d, Video S4). Inter-subunit β-sheets shift downward in concert with these rotations: CCT3–CCT1 by 1.9 Å, CCT1–CCT4 by 2.5 Å, CCT5–CCT7 by 4.1 Å, and CCT7–CCT8 by 1.8 Å, corresponding to the equatorial domain rotations of CCT1, CCT5, and CCT7 (Figure 3j). Other inter-subunit β-sheets undergo only minor adjustments (Figures S14a, S14b).

From R1 to R5, although some of the same type of subunits between the two rings tilt differently in their apical domains (Figure S8), the conformational changes in the intermediate and equatorial domains remain similar. In each state from R6 to R8, the two rings display high structural similarity; therefore, C2 symmetry was applied during reconstruction. In state R6, resolved at 3.8 Å, the protrusion network further stabilizes CCT8 and now extends to include CCT1 (Figures 1, 3b, Figures S6, S7, S8, Video S5). Beneath the protrusion network, the apical loops of CCT2, CCT4, CCT5, CCT7, and CCT8 assemble into a β-sheet, reflecting ongoing formation of the β-sheet barrel, supported by the improved density with better resolved density of β strands in this region (Figure 3d). While the protrusion helix of CCT1 anchors towards the apex, its apical loop has not yet contributed to β-sheet formation. CCT3 remains adjacent to CCT1, creating a substantial gap between CCT3 and CCT6 (Figure 1).

ATP is bound in all subunits, and a water molecule is now observed adjacent to the ATP in every subunit except CCT3, which remains relatively open (Figure 3c, Figure S10). From R5 to R6, all subunits undergo further equatorial domain rotations, with CCT1 exhibiting the largest change—an additional 9.5° rotation within the ring (Figures 3g, 3i, Figures S13e, S14a). Corresponding β-sheet shifts are observed: CCT7–CCT8 by 1.1 Å, CCT6–CCT3 by 1.5 Å, CCT3–CCT1 by 2.4 Å, and CCT1–CCT4 by 1.7 Å, while other interfaces show only minor displacements (Figure 3j, Figure S14b, Video S5).

Relative to R5, in R6 the intermediate domain of CCT1 rotates downward by 31.5° toward the equatorial domain, nearly closing its nucleotide-binding pocket (Figure 3h, Figure S12). The apical domain rotates in concert with the intermediate domain, producing an inward tilt of the protrusion helix tip of approximately 51 Å (Figure S13e). From R1 to R5, CCT3 engages CCT1 via their apical domains, using complementary electrostatic surfaces formed by helices and flexible loops (Figure 3l). Although CCT1 has been attempting the closure, it is blocked by CCT3 (Videos S1, S2, S3, S4). In state R6, the apical domain of CCT3 has repositioned away from the CCT1 closure trajectory, resolving the steric interference observed in R1-R5 and allowing CCT1 to proceed with closure (Figure 3k). CCT3, in contrast, displays lower resolvability in its apical domain (Figure 1, Figure S7). Unlike previous states, where CCT3’s apical domain tilted toward CCT4, in R6 it tilts back toward alignment with its equatorial domain. From R5 to R6, the apical domain of CCT3 rotates ∼23° relative to the intermediate domain, producing an estimated inward tilt of the protrusion helix tip of ∼40 Å, though this value is approximate due to limited resolution (Figure S13e).

CCT1 acts as the primary responder to ATP binding in R6. Its incorporation into the protrusion network during the R5→R6 transition is driven by ATP-induced equatorial rotation and apical tilting, with the equatorial rotation likely aided by inter-ring cooperativity via CCT7′ in the opposite ring. The closure of CCT1, following CCT7 closure in R5, further destabilizes the support base formed by CCT1, CCT3, CCT6, CCT8, and CCT7 in earlier states, promoting equatorial domain rotations across the remaining subunits (Figure 3a, 3f, 3g, Figure S14a, Video S5).

### The Simultaneous but Distinct Closure of Hemisphere CCTs 8, 6, 3

In state R7, resolved at 3.9 Å, CCT1, CCT4, CCT2, CCT5, CCT7, and CCT8 form a robust protrusion network (Figure 1, Figure 4b, Figure S6, S7, S8, Video S6), with their apical loops assembling into a continuous β-sheet beneath the protrusions, further extending the β-sheet into a barrel relative to R6 (Figure 4d). The protrusions of CCT6 and CCT3 appear to be joining this network, although their weak density suggests considerable structural flexibility (Figure 1, 4a). Notably, the previously large gap between CCT6 and CCT3 is narrowing.

**Figure 4.**
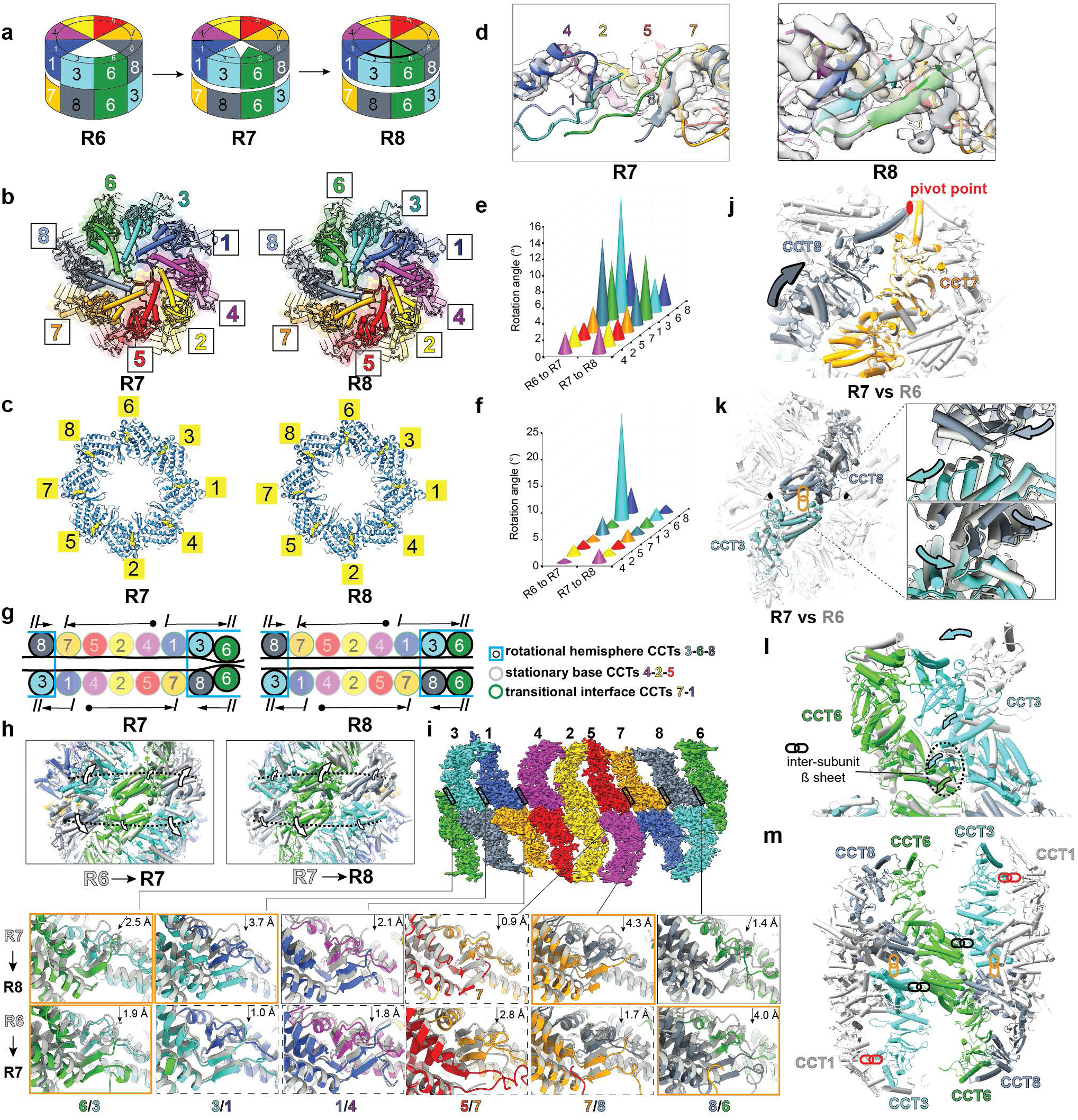
The closure of CCTs 3-6-8. **a.** The schematic of sequential closure of CCTs 3-6-8 in R7 and R8. **b**. The top view of structural models of R7 and R8, showing the sequential extension of apical protrusion network by CCTs 3-6. **c**. The ATPMg^2+^H2O (yellow) distribution in R7, R8. **d.** The extension and completion of apical β-barrel in R7 and R8. **e**. The histogram quantifying the equatorial domain rotation across R6, R7 and R8. **f.** The histogram quantifying intermediate domain rotation towards equatorial domain across R6, R7 and R8. **g.** The unwrapped arrangement of CCTs in TRiC in states R7 and R8, demonstrating the sequential response to ATP binding by CCT3-8 and CCT6. **h**. The inter-ring separation by equatorial rotation at CCT3-6-8 across R6, R7, R8. **i.** The inter-subunit betasheet transitting the allostery across R6, R7, R8. **j**. CCT8 rotates around its protrusion tip as pivot point from R6 to R7. **k.** The concerted equatorial rotation of CCT3 and CCT8 from R6 to R7. **I.** The overlap of R1 (gray) and R7 (colored) showing the restraint on CCT6 by the inter-subunit β sheet from CCT3 is released in R7. **m.** The allosteric restaints by CCT3 on CCT8, CCT6, CCT1.

All CCT subunits in R7 contain ATP in their nucleotide-binding pockets, with a water molecule adjacent to each ATP (Figure 4c, Figure S10). Compared to R6, the intermediate domain of CCT1 rotates downward toward its equatorial domain by 4.8°, further closing its nucleotide-binding pocket (Figure S12). CCT3 rotates its intermediate domain downward by 20.1° and aligns its apical and intermediate domains with the equatorial domain, adopting the same overall domain arrangement as the other subunits (Figure 4f). This alignment enables CCT3’s protrusion helix to approach and partially join the protrusion network.

Relative to R6, all CCT subunits undergo further equatorial domain rotations (Figures 4e, 4h, S13f, S14a). CCT3 and CCT8 show the largest changes (14.4° and 10.8°, respectively), while CCT6 rotates substantially by 7.2°. CCT1 and CCT7 have smaller rotations (6.6° and 3.4°), whereas CCT4, CCT2, and CCT5 remain largely stationary. Notably, from R6 to R7, the hemisphere comprising CCT4, CCT2, and CCT5 switches roles to form a stable support base, allowing the remaining subunits—CCT1, CCT3, CCT6, CCT7, and CCT8—to rotate around it (Figure 4g, Video S6). Corresponding inter-subunit β-sheet movements reflect these rotations: downward shifts occur between CCT7–CCT8 (4.3 Å), CCT3–CCT1 (3.7 Å), and CCT1–CCT4 (2.1 Å), while lateral shifts are seen between CCT8–CCT6 (1.4 Å) and CCT6–CCT3 (2.5 Å) (Figure 4i). In contrast, β-sheets connecting the support-base subunits—CCT4–CCT2, CCT2–CCT5, and CCT5–CCT7—exhibit less movement (Figure 14b).

Across R1–R7, the intermediate domains of CCT6 and CCT8 remain fixed relative to their equatorial domains, with helix 12 positioned as in the fully closed TRiC state (R8, discussed later) (Figure S12). In earlier states, the movement of CCT8’ was restricted by CCT3 in the opposing ring; in R7, however, CCT3’s equatorial rotation enables substantial rotation of CCT8’ through inter-ring cooperativity (Figure 4k, 4m). CCT8 also undergoes ∼–23° rotation of its apical domain relative to the intermediate domain (Figure S15a). Early incorporation into the protrusion network in R5 positions the tip of its protrusion helix as a pivot point, enabling the apical domain to rotate in R7 with minimal displacement at the tip (Figure 4j). Similarly, CCT6 —located on the C2 symmetry axis between two rings—is constrained by intra-ring interactions with CCT3 in states preceding R7 (Figure 4l, 4m). This constraint is released during the R6–R7 transition, as evidenced by lateral shifts in inter-subunit β-sheets connecting CCT8–CCT6 and CCT6–CCT3 (Figure 4i).

State R8, resolved at 3.5 Å, represents the fully closed TRiC conformation, with all eight CCT subunits closed, and a complete protrusion network formed (Figure 1, 4b, Figure S6, S7, S8, Video S7). The apical loops assemble into a compact β-sheet barrel at each chamber apex (Figure 4d). ATP is bound in every nucleotide-binding pocket, with a water molecule adjacent to each ATP (Figure 4c, Figures S10 and S11). Compared to R7, all CCT subunits undergo additional equatorial rotations (Figure 4e, 4h, S13g, S14a). CCT6 rotates most prominently (8°), followed by CCT8 and CCT3, which rotate 5.2° and 4.2°, respectively, while the remaining subunits exhibit more modest rotations (Figure 4e, 4h, Video S7). Significant downward shifts occur in inter-subunit β-sheets between CCT5–CCT7 (2.8 Å), CCT7–CCT8 (1.7 Å), CCT8–CCT6 (4.0 Å), CCT6–CCT3 (1.9 Å), and CCT1–CCT4 (1.8 Å), consolidating the closed architecture (Figure 4i).

Intermediate-to-equatorial domain alignment remains largely unchanged from R7 to R8, though CCT4, CCT6, and CCT7 adjust their apical domains to better conform to the chamber wall (Figure S12, S15b). Overall, R8 adopts a more compact conformation than the ADP/AlFx-locked TRiC structure (PDB:7TTT)^7^, with a lower-positioned β-sheet barrel at the chamber apex (Figure S15c). Helix 12 of each intermediate domain, containing the conserved aspartic acid and coordinated water, is closer to the nucleotide-binding pocket than in the AlFx-locked structure. ATP shifts toward the coordinated water, highlighting a key structural distinction between ATP binding and ADP/AlFx locking (Figure S15c). TRiC likely requires an even more compact closed conformation than R8 so that the water molecule can be delivered closer to the γ-phosphate for a nucleophilic attack.

In summary, CCT3 is the primary responder to ATP binding from R6 to R7, relieving inter-ring constraints on CCT8’ and intra-ring constraints on CCT6, which promotes substantial equatorial rotations in both subunits (Figure 4m, Video S6). In R8, all CCT subunits continue equatorial rotations, most prominently in CCT6, CCT8, and CCT3, while CCT4, CCT6, and CCT7 flexibly adjust their apical domains to conform to the compact chamber (Video S7). Notably, ATP binding alone drives TRiC into a more closed conformation than the ADP/AlFx-locked state.

## Discussion

### Mechanism of Sequential and Asymmetric Ring Closure in Human TRiC

The chaperonin TRiC facilitates folding of the eukaryotic cytosolic proteome through an ATP-driven cycle of chamber closure. This study dissects at near atomic resolution the temporal sequence of subunit engagement during human TRiC ring closure triggered by ATP binding. ATP binding—not hydrolysis—couples protrusion network extension and equatorial domain rotation to sequentially close the ring in human TRiC (Figure 5a, 5b, 5c). The closure process begins at CCT4, the first subunit to anchor its protrusion helix near the central vertical axis of the chamber, and TRiC progressively extends the protrusion network in the order: CCTs 4 > 2 > 5 > 7 > 8 > 1 > 3/6. This network forms through differential response to ATP binding in TRiC subunits with intra-ring equatorial domain rotations, following a distinct order: CCTs (4 > 2 > 5) > 7 > 1 > (3/8 > 6) (Figure 5b). The observed sequences reflect a hierarchy of conformational responses at each step of the transition, including ATP-driven subunit-specific responses, local intra-ring cooperativity and inter-ring allosteric coupling, rather than a simple intra-ring nearest-neighbor transmission pathway.

**Figure 5.**
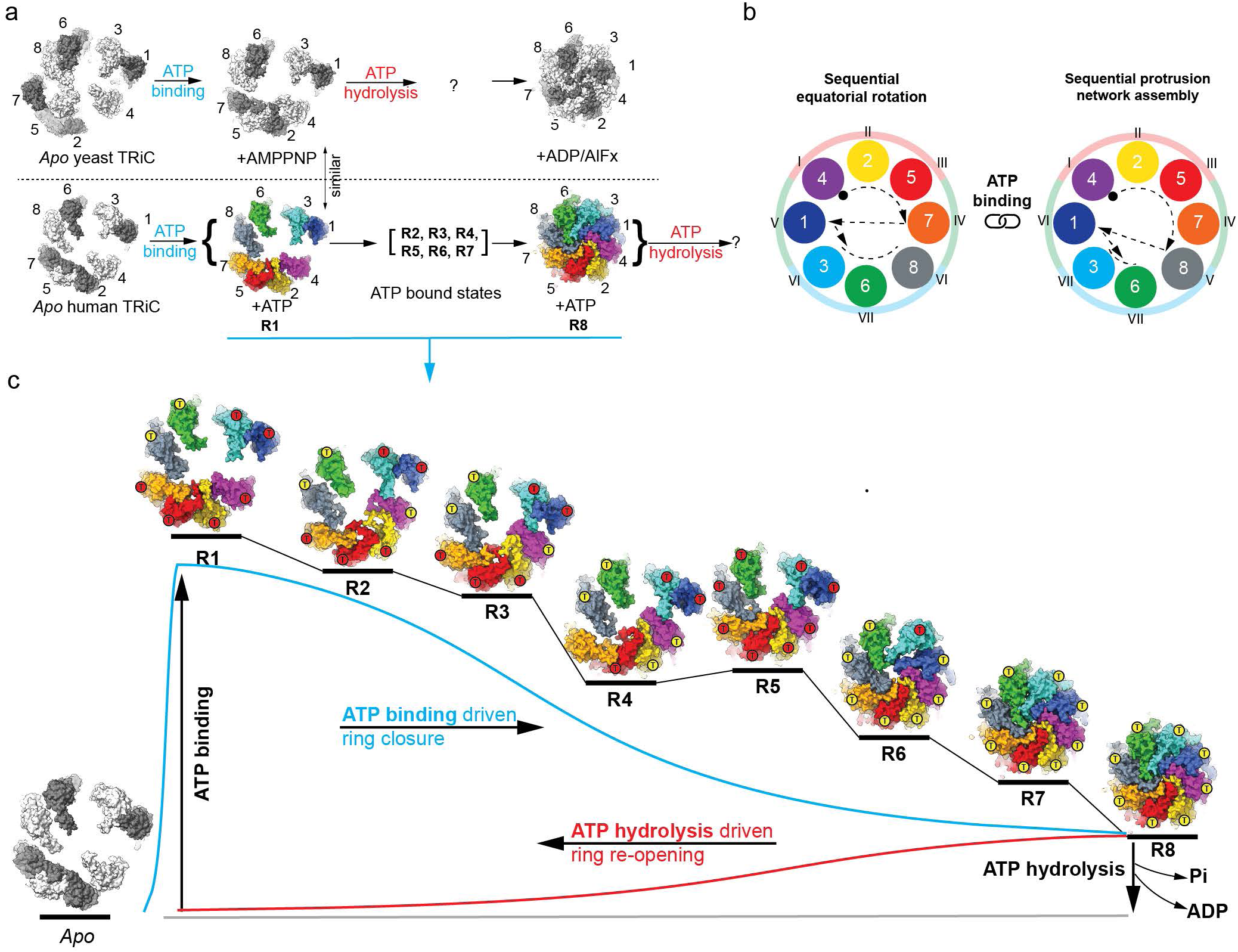
The ATP binding driven allosteric scheme of sequential ring closure in human chaperonin TRiC. **a.** Top panel: The current understanding of yeast TRiC ring closure with ATP analogue. Bottom panel: The observed human TRiC ring closure triggered by ATP binding in this study, **b**. ATP binding coupling sequential apical protrusion network extension and equatorial domain rotation in human TRiC ring closure, **c.** The eight observed TRiC-ATP states mapped onto a conformational landscape. The landscape shows TRiC conformational free energy as a function of ring-closure progression, conditioned on ATP-occupancy states measured under our experimental conditions (1 μM TRiC, 50 μM ATP, 20 s incubation). Because the observed R4 state lacks two ATP molecules (one in each CCT7 subunit), we place R4 in a lower local energy minimum than R5, since progression from this ATP-deficient R4 to R5 would be energetically unfavorable. Conversely, a hypothetical R4 with ATP bound in all CCT subunits is expected to occupy a higher-energy minimum than R5 and thus promote progressive closure.

In addition to the differential response to ATP binding among subunits, human TRiC uses intrinsic apical domain interactions to enforce the asymmetric ring closure. CCT7 forms apical contacts with CCT5, likely preventing premature inward tilting of CCT5’s protrusion helix and regulates the sequential closure at the CCTs 4-2-5 hemisphere. CCT7 with lowest ATP-binding affinity, dissociates from CCT5 in the absence of ATP, but upon ATP binding, its closure recruits CCT8 into the protrusion helix network. A second apical interaction between CCT3 and CCT1 delays the closure of the CCT1-3-6-8 hemisphere: CCT3 sterically hinders CCT1 with its apical domain, prevents the equatorial rotation of CCT8 in the opposite ring via inter-ring interface, and restrains CCT6 with the inter-subunit intra-ring β-sheets. Disruption of the CCT3–CCT1 contact allows CCT3 to tilt toward CCT6, closing the large gap between them. This repositioning relieves intra-ring constraints on CCT6 and inter-ring constraints on CCT8, enabling coordinated equatorial domain rotation and triggering closure of the CCT3-6-8 hemisphere.

The pre-closure apical interactions between CCT5–CCT7 and CCT1–CCT3 act as inhibitory checkpoints that enforce sequential, asymmetric ring closure. CCT7 and CCT1 serve as transitional interfaces between the CCT4–2–5 and CCT3–6–8 hemispheres during the closure process. Closure begins with equatorial rotation of the CCT4–2–5 hemisphere while CCT3–6–8 remains stationary. The motion propagates through CCT7 and CCT1 to CCT3–6–8, at which point roles reverse: CCT3–6–8 rotates while CCT4–2–5 stabilizes the structure (Figure 5b).

Importantly, ATP loading occurs faster than conformational closure: all subunits can bind ATP within seconds, yet closure proceeds sequentially over the 20-second timescale. This suggests that closure order is governed by conformational coupling rather than ATP-binding kinetics alone. Even with all sites occupied, structural constraints within TRiC play a significant role in determining which subunits close first. Thus, ATP binding alone does not drive immediate subunit closure; instead, a sequential release of apical allosteric restraints permits complete ring closure. These findings indicate that the relationship between ATP affinity and closure timing is more complex than a simple 1:1 correspondence and involve species-specific differences and allosteric regulation.

Previous studies reported divergent nucleotide-binding gradients among the eight CCT subunits of bovine and yeast TRiC. The crosslinking study of bovine TRiC with 8-azido-ATP reported that CCT5 and CCT4 showed the highest affinities, followed by CCT1 and CCT2. In contrast, CCT3, CCT6, CCT7, and CCT8 exhibited very low ATP affinity and, even at high ATP concentrations, displayed minimal ATP occupancy^16^. In contrast, a cryo-EM study of yeast TRiC reported a different nucleotide-affinity gradient using ADP·AlFx: highest affinity in CCT3, CCT6, and CCT8; intermediate affinity in CCT1 and CCT7; lower affinity in CCT2 and CCT5; the lowest affinity in CCT4^17^, and a high ADP affinity for CCT8^14^. Our analysis of human TRiC reveals only CCT7 as the lowest-affinity subunit; the ATP-affinity hierarchy among the other subunits remains unresolved. Whether human TRiC exhibits a similar ATP-affinity gradient to that reported for bovine or yeast TRiC, and whether subunit closure order correlates with such a gradient, remains to be investigated. Our data shows that CCT7, characterized as low-affinity in bovine TRiC but intermediate in yeast, undergoes closure relatively early, while CCT1, reported as high-affinity in bovine TRiC but intermediate in yeast, closes relatively late. The extent to which ATP binding affinity hierarchy in human TRiC regulates ring closure remains unknown and is an important subject for future study.

### Energy-Efficient TRiC Ring Closure Evolved for Substrate-Specific Folding

Our work reveals the mechanism of human TRiC ring closure and provides critical insights into a process that has remained poorly understood in eukaryotes. In sharp contrast to the current model based on yeast or bovine TRiC studied with a nonhydrolyzable ATP analogue^14,15^—which suggested that ATP hydrolysis is required for ring closure—our findings demonstrate that ATP binding alone is sufficient to drive closure (Figure 5a). TRiC delays ring closure through allosteric restraints between apical domains and the low ATP affinity of CCT7, and ATP hydrolysis proceeds only once the ring is fully closed. These features work together to ensure efficient energy use and prevent futile ATP hydrolysis in partially closed rings.

We observe full ATP occupancy in the majority of TRiC particles at 50 μM ATP, in stark contrast to a prior report using 8-azido-ATP crosslinking that detected only 8 ATPs per bovine TRiC complex even at much higher ATP concentrations^16^. Based on our ATP-bound structures, 8-azido-ATP crosslinking may detect only crosslink-competent subunits and thus underestimate overall ATP binding. Our structures capture TRiC during ATP-driven ring closure, representing the physiologically relevant forward pathway of ring closure. However, no structural study to date has captured TRiC exactly at the moment of ATP hydrolysis or in *bona fide* ADP-bound, native post-hydrolysis conformations. Previous single-molecule measurements showing retention of 8 ADP molecules per bovine TRiC after ATP incubation and nucleotide washout can be explained by differential ADP release kinetics across TRiC subunits, rather than a model in which only eight ATPs are hydrolyzed per cycle^12^. Another study examined yeast TRiC in the presence of high concentrations of exogenous ADP^18^, but it remains unclear whether those conditions recapitulate native post-hydrolysis states or instead induce alternative conformations. In the native catalytic cycle, ATP binding drives ring closure (as shown here), followed by hydrolysis and ADP/Pi release that drive reopening. Post-hydrolysis intermediates, potentially comprising mixed ATP/ADP occupancy, Pi release in progress, and partially reopened rings, are likely short-lived and structurally heterogeneous. Definitive characterization of the conformational pathway during and after hydrolysis will require resolving authentic post-hydrolysis ensembles that arise from ATP turnover within closed TRiC, rather than from addition of exogenous ADP to apo-TRiC.

Unlike homologous bacterial chaperonins such as GroEL, whose subunits within a ring close simultaneously, the hetero-oligomeric TRiC undergoes asymmetric, sequential ring closure. This stepwise mechanism may have evolved to accommodate a more complex spectrum of TRiC client proteins and their variable abundances, enabling the folding of certain eukaryotic proteins that GroEL cannot handle^2,19,20^. By coordinating substrate folding with the kinetics of ring closure, as exemplified by the segmental release of actin^11^, TRiC ensures efficient folding within its natural timing. In our study, human TRiC was purified from insect cells overexpressing the complex (Figure S16a), with no exogenous client proteins added prior to cryo-EM experiments. Although particles of R1-R8 appear client-free by the latent space analysis with CCT tail density at the inter-ring cavity (Figure S16b, Movies S8 and S9), we cannot exclude the possibility that a small fraction in the full dataset retained endogenous clients (e.g., tubulin), which might have slightly affected the kinetics or the equilibrium distribution of conformational states. Despite this limitation, our study establishes a baseline ATP-binding–induced ring-closure pathway and lays the groundwork for future investigation of client-specific effects. Future studies could explore how the timing of subunit closure and substrate-specific interactions are fine-tuned across different clients, providing deeper insight into the evolutionary and mechanistic principles underlying TRiC’s folding versatility.

### Functional and Therapeutic Implications of TRiC Ring Closure

The sequential closure of the TRiC ring is likely tied to efficient substrate capture. The delayed closure of the CCT3–6–8 hemisphere may allow extra time for recruiting substrates—such as tubulin^7,10^ or actin intermediates^6^—that preferentially associate with these subunits. A rat study showed that a CCT4 (C450Y) missense mutation causes hereditary sensory neuropathy, with sensory neuron loss and severe ataxia^21,22^.In humans, missense mutations in CCT1 (K159R) and CCT5 (K176R) within the nucleotide-sensing loop are associated with brain malformations and seizures. These substitutions likely disrupt nucleotide sensing in the affected subunits, leading to allosteric defects in TRiC that undermine microtubule and actin cytoskeletal organization^23^.

Understanding the TRiC ring closure mechanism is crucial for drug design because this ATP-driven conformational change governs how TRiC encapsulates and folds client proteins such as actin^6,11,24^, tubulin^7,10^, the oncogenic protein STAT3^25^, and tumor suppressor protein p53^26^. Many diseases, including Huntington’s^27–30^, Alzheimer’s^31,32^, and Parkinson’s^33,34^, involve protein misfolding and aggregation, and TRiC has been shown to interact with and modulate aggregation-prone proteins such as huntingtin and α-synuclein. Structural insights into this process uncover allosteric sites and ATP-binding pockets that could be exploited to modulate TRiC activity. Compounds that stabilize particular conformations, alter ATPase-driven closure, or mimic features of the folding chamber could selectively boost or suppress its function. In this way, our delineation of the molecular basis of TRiC ring closure lays the groundwork for developing therapeutics that fine-tune TRiC activity and restore proteostasis in disease.

## Materials Availability

Reagents generated in this study are available from the lead contact upon request without restriction.

## Data and Code Availability

The model and maps are accessible in EMDB and PDB (EMD-70534, EMD-70535, EMD-70537, EMD-70538, EMD-70539, EMD-70540, EMD-70541, EMD-70542, 9OSQ, 9OSP, 9OSJ, 9OSH, 9OSE, 9OS9, 9ORU, 9ORR). No code is generated in this manuscript. Further information and requests for resources and reagents should be directed to Yanyan Zhao Any additional information required to reanalyze the data reported in this paper is available from the lead contact upon request.

## Acknowledgements

We acknowledge funding support from the Stanford School of Medicine Dean’s Postdoctoral Fellowship (to Y.Z.). The image data was collected at Stanford-SLAC CryoEM Center (S^2^C^2^) which is supported by the National Institutes of Health Common Fund Transformative High-Resolution Cryo-Electron Microscopy Program (R24GM154186). We thank Dr. Judith Frydman for her support, Dr. Muyuan Chen for advice on data presentation, and Grigore Pintilie for guidance on Q-score interpretation.

## Author contributions

Conceptualization: Y.Z.; Methodology: Y.Z.; Investigation: Y.Z.; Writing – original draft: Y.Z.; Data interpretation, review & manuscript editing: Y.Z., M.S., W.C.; Funding acquisition: Y.Z., W.C..

## Method

### Protein overexpression and purification

A GFP tag was inserted after CCT1 residue E479 in the equatorial domain, with short linkers flanking GFP, to label the human TRiC complex and facilitate CCT subunit identification (Figure S5). Human TRiC complex was expressed using the Bac-to-Bac baculovirus system (Invitrogen). Hi5 cells (1 L culture) were infected with P3 baculovirus encoding the GFP-tagged TRiC complex. Infected cells were incubated at 27°C and harvested after ∼72 hours. Cells were resuspended in the lysis buffer (20 mM HEPES pH 7.4, 50 mM NaCl, 10% glycerol, 5 mM PMSF, 1 mM DTT) supplemented with DNase I and protease inhibitors. Lysis was performed using an Emulsiflex, and cell debris was removed by centrifugation at 4°C for 20 minutes. TRiC was precipitated from the supernatant using ammonium sulfate and collected by centrifugation at 20,000 × g. The pellet was resuspended and dialyzed overnight in the same buffer. The dialyzed sample was applied to a heparin column and eluted with a linear gradient up to 1.0 M NaCl. TRiC-containing fractions were concentrated and further purified by size-exclusion chromatography on a Superose 6 10/300 GL column. Eluted fractions were analyzed by SDS-PAGE, and TRiC-positive fractions were pooled and concentrated.

### Cryo-EM sample preparation and data collection

ATP was added to purified TRiC in ATPase buffer (30 mM Tris-HCl, pH = 7.4, 100 mM KCl, 5 mM MgCl_2_ and 1 mM DTT) to initiate the reaction (at t = 0 s) with a final concentration of 1 μM TRiC and 50 μM ATP, and 3 μL of the sample was applied to glow-discharged Quantifoil R2/1 grids, blotted for 3 seconds at 21 °C, and plunge-frozen in liquid ethane (at a time point of approximately 20 s). Grids were then kept in liquid nitrogen and loaded into a Titan Krios G3i cryo-electron microscope operated at 300 kV. Data collection was performed using a Falcon 4i camera at a pixel size of 1 Å, with a total electron dose of 49.63 e⁻/Å². A total of 24,287 movie stacks were acquired, each consisting of 40 frames. The defocus range during data collection was set from −1.5 μm to −2.5 μm.

### Data preprocessing

Movie stacks were aligned and dose-weighted using MotionCor2^35^. In RELION v3.1^36^, TRiC particles were automatically picked using a template-matching approach with a box size of 320 × 320 pixels. Particles were refined through iterative rounds of 2D and 3D classification to remove false positives. A total of 1,630,427 particles contributed to the reconstruction of the open-state TRiC at 3.53 Å resolution, with no symmetry imposed. An additional 221,070 particles were used to generate a closed-state TRiC reconstruction at 3.45 Å resolution, with C2 symmetry applied.

### Heterogeneity analysis

For each particle set including the open and closed states, images were initially downsampled to 80 × 80 pixels (4 Å/pixel) and input into a cryoDRGN v2.0^37^ neural network for heterogeneity analysis, featuring an 8-dimensional latent space and encoder/decoder architectures of size 1024 × 3. After 50 training epochs, 200 volumes were generated at the k-means cluster centers of the latent embeddings and visualized in ChimeraX to identify and discard low-quality (“garbage”) particles. The remaining particles were then downsampled to 160 × 160 pixels (2 Å/pixel) and reprocessed using the same neural network architecture in multiple iterative rounds to classify conformational heterogeneity.

In each round, 3D volumes were again generated at the cluster centers of the latent space and examined in ChimeraX^38^. Particles corresponding to similar conformations were grouped and used for subsequent rounds of cryoDRGN^37^ training and analysis. This iterative heterogeneity analysis continued until convergence, defined as a minimal presence of minority conformations in each run. Ultimately, particles corresponding to each of the eight conformational classes identified in this study were extracted and reconstructed by local auto-refinement in RELION v4.0^39^, using the average reconstructions of the open- and closed-state reconstructions above as the references. During reconstruction of states R6, R7, and R8, C2 symmetry was imposed between the two rings (with the symmetry axis passing through CCT2 and CCT6) to improve resolution, as symmetry-free reconstructions demonstrated that the two rings adopt nearly identical conformations in these closed states. This symmetry imposition does not affect subunit assignment, which was confirmed by the visualization of the GFP tag at the equatorial domain of each CCT1 subunit. For states R1–R5, no symmetry was imposed, preserving the asymmetric features and clear GFP density.

### Model building

For all reconstructions, modeling of CCT subunits followed a hierarchical and resolution - dependent strategy. Each CCT subunit model of the closed-form TRiC complex (PDB: 7TTT) was segmented into three structural domains: the equatorial, intermediate, and apical domains. For each state, the domains of each subunit were individually fitted into the corresponding unsharpened map using rigid-body docking to determine their overall orientations at intermediate resolution. Subsequent local refinement against sharpened maps was performed using ISOLDE^40^ to improve local geometry and fit, but only where the local resolution justified it (6-8 Å for helical features, 4.5-4.7 Å for β-strands, and ∼3.5 Å or better for residue assignment). In regions where the sharpened maps were noisy or the local resolution was insufficient—most notably in the apical domains—no secondary-structure or side-chain refinement was carried out. All figures were generated using ChimeraX^38^. The resolvability of ATP and water in the nucleotide binding pocket is quantified by Q score^41,42^ using MapQ from Segger in Chimera v1.18.

